# Trigeminal nerve direct current stimulation causes sustained increase in neural activity in the rat hippocampus

**DOI:** 10.1101/2023.12.12.571341

**Authors:** Liyi Chen, Zhengdao Deng, Boateng Asamoah, Myles Mc Laughlin

## Abstract

Transcranial direct current stimulation (tDCS) is a noninvasive neuromodulation method that can modulate many brain functions including learning and memory. Recent evidence suggests that tDCS memory effects may be caused by co-stimulation of scalp nerves such as the trigeminal nerve (TN), and not the electric field in the brain. The TN gives input to brainstem nuclei, including the locus coeruleus that controls noradrenaline release across brain regions, including hippocampus. However, the effects of TN direct current stimulation (TN-DCS) are currently not well understood. In this study we hypothesized that TN-DCS manipulates hippocampal activity via an LC-noradrenergic bottom-up pathway. We recorded neural activity in rat hippocampus using multichannel silicon probes. We applied 3 minutes of 0.25 mA or 1 mA TN-DCS, monitored hippocampal activity for up to 1 hour and calculated spikes-rate and spike-field coherence metrics. Subcutaneous injections of xylocaine were used to block TN and intraperitoneal injection of clonidine to block the LC pathway. We found that 1 mA TN-DCS caused a significant increase in hippocampal spike-rate lasting 45 minutes in addition to significant changes in spike-field coherence, while 0.25 mA TN-DCS did not. TN blockage prevented spike-rate increases, confirming effects were not caused by the electric field in the brain. When 1 mA TN-DCS was delivered during clonidine blockage no increase in spike-rate was observed, suggesting an important role for the LC-noradrenergic pathway. These results provide a neural basis to support a tDCS TN co-stimulation mechanism. TN-DCS emerges as an important tool to potentially modulate learning and memory.

**Highlights:**

1. Trigeminal nerve direct current stimulation (TN-DCS) boosts hippocampal spike rates
2. TN-DCS alters spike-field coherence in theta and gamma bands across the hippocampus.
3. Blockade experiments indicate that TN-DCS modulated hippocampal activity via the LC-noradrenergic pathway.
4. TN-DCS emerges as a potential tool for memory manipulation.

**Figure Graphic Abstract:** 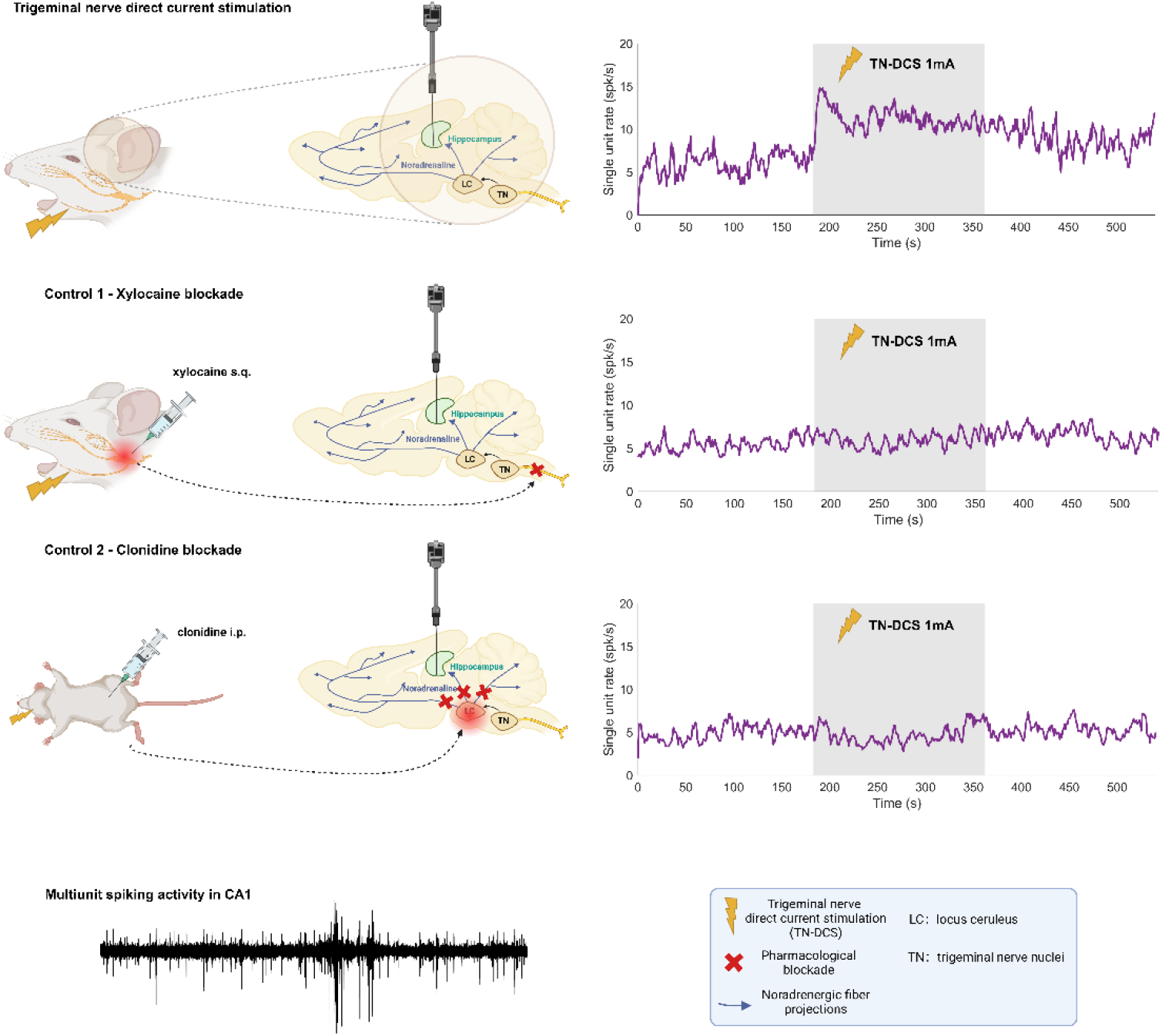

## Introduction

Transcranial direct current stimulation (tDCS) is a popular noninvasive brain stimulation method that has been used to modulate a wide range of brain functions in both healthy volunteers and patients [1–3]. In healthy volunteers tDCS has been shown to modulate motor memory [4], working memory [5] and re-consolidation of long-term memory [6]. While in Alzheimer’s disease tDCS can improve both recognition memory [7] and cognitive function [8]. tDCS works by passing an electrical current of 1-2 mA through scalp electrodes to create a weak electric field in the brain of typically less than 1 V/m [9–11]. The electric field in the brain polarizes the membrane potential in cortical neurons, that in turn modulates their excitability [12]. This effect on cortical regions may propagate via a top-down pathway to other interconnected brain regions such as the hippocampus, influencing functional networks, and synchronization between these regions [13]. This direct mechanism is generally assumed to underly the observed tDCS effects. However, the precise tDCS mechanism (or more likely mechanisms) remains poorly understood and is still the subject of ongoing research and debate within the field [14].

A number of recent studies have put forward a different hypothesis of how tDCS may work [15–17]. In addition to the weak electric field in the brain, tDCS also generates a stronger electric field in the scalp of up to 20 V/m [18, 19]. This field is strong enough to stimulate cranial and cervical nerves [20] in the scalp such as the greater occipital and trigeminal nerves. Both the trigeminal and occipital nerves give direct input to the tactile circuit (e.g. thalamus and somatosensory cortex [21]) which accounts for the tingling and itching sensations reported by tDCS subjects [22]. Furthermore, these nerves give input to a series of brain-stem nuclei including the solitary tract nucleus and the locus coeruleus (LC) [23]. The LC is the key nucleus in the noradrenergic system which projects to many different brain regions, including strong projections to the hippocampus [24, 25]. This noradrenergic mediated LC-hippocampus connection can regulate neural activity, effect neural plasticity [26] and modulate memory performance [27]. Recent evidence found that noradrenergic fibers originating from the LC were the main source of neurotransmitter acting on dopaminergic receptors in the dorsal hippocampus [28, 29] - termed the LC-dopaminergic pathway. Noradrenergic stimulation/suppression causes a parallel change in concentration of both noradrenaline and dopamine [30]. Thus, there is a bottom-up pathway from the tDCS site on the scalp, via the trigeminal and occipital nerves, to the LC-noradrenergic system and then the hippocampus. This bottom-up pathway could account for some tDCS-memory effects [15]. However, the potential contribution of a tDCS bottom-up pathway has only recently begun to be investigated.

A study by Vanneste et al in [17] was the first to show that tDCS effects on memory may be caused by indirect stimulation of the greater occipital nerve. A series of studies in both healthy volunteers and rats indicated that these effects were mediated via the LC-noradrenergic system. A follow-up randomized controlled clinical trial from the same group suggested that DC stimulation of the greater occipital nerve could boost associative memory in older adults[31]. In a final study the same group showed that DC stimulation of the greater occipital nerve can boost long-term memory retention and presented evidence suggesting this effect was mediated via a greater occipital nerve LC-dopaminergic pathway to the hippocampus [32].

Theta and gamma oscillations are known to play a central role in hippocampal memory formation [33]. Of particular importance for memory circuits is the coherence between individual neurons (spikes) and the specific local field potential oscillation (LFP) [34, 35]. Higher spike-field coherence (SFC) suggests that the neurons are firing more consistently and precisely in relation to the ongoing oscillation, usually in the theta (4-8 Hz) or gamma (30-100 Hz) frequency range. In the context of memory and learning, increased SFC is associated with improved neural communication and more effective neural coding [35, 36] which plays a crucial role in memory and cognition [37]. Importantly, dopamine and noradrenaline concentrations are associated with changes in hippocampal excitability and SFC [38–40]

In the most standard tDCS montage the cathode is placed in a supraorbital position while the anode is placed above the motor cortex [3]. This montage stimulates both the trigeminal and the greater occipital nerves. In this study, we tested the hypothesis that stimulation of the trigeminal nerve with direct current (TN-DCS) would alter neural hippocampal activity. Our results showed that TN-DCS in the rat significantly increased neural activity in the hippocampus with effects lasting up to 45 minutes. Blocker experiments indicated these effects were caused by LC-noradrenergic pathway activated via TN stimulation. Additionally, TN-DCS significantly altered theta and gamma band SFC in the hippocampus.

## Material and methods

### Animals

7 male Sprague-Dawley rats (250-400 grams, Charles-river laboratories) were housed in a colony (19°C, 14/10-hour light/dark cycle) and had unrestricted access to food and water. All procedures were approved by the KU Leuven ethics committee for laboratory experimentation (P072/2020).

### Surgery and preparation

Rats were anesthetized (intraperitoneal urethane, 10 mg/mL, Sigmaaldrich, USA) and placed in a stereotaxic frame (Narishige type SR-6, No. 7905), and their core temperature monitored via a metal rectal probe. Anesthesia level was routinely monitored using the toe-pinch reflex. We exposed the skull and corrected the stereotaxic positioning by ensuring the dorso-ventral offset between bregma and lambda did not exceed 0.2 micrometers. We then drilled a burr hole (US#4 HP-014 drill bit, Meissinger, Germany) relative to bregma at 3.45mm AP, 2.35mm ML to target the right dorsal hippocampus.

### Electrophysiological recording setup

For electrophysiology recordings we used a one column, 32-channel single probe spanning 1550 μm (E32+R-50-S1M-L20 NT, Atlas Neuro, Leuven, Belgium) with a pointy tip. Signals from the probe were amplified (×192), bandpass filtered (0.1 Hz to 7.9 kHz), and digitized (16 bit, 30 kHz) using an RHD 32 Intan head-stage (Intan Technologies, Los Angeles, CA) and an Open-Ephys acquisition board (www.open-ephys.org). The digitized signals were visualized and stored on a PC hard-drive using the Open-Ephys GUI v0.6.4. The probe was inserted into the hippocampus through the craniotomy and spanned 3400 (deepest) to 1850 μm (uppermost electrode). Based on depth, multi-unit activity and the Paxinos atlas [41] we assigned putative hippocampal layers (CA1, DG, CA3) to different electrode channels (Fig. 1).

**Figure 1.**
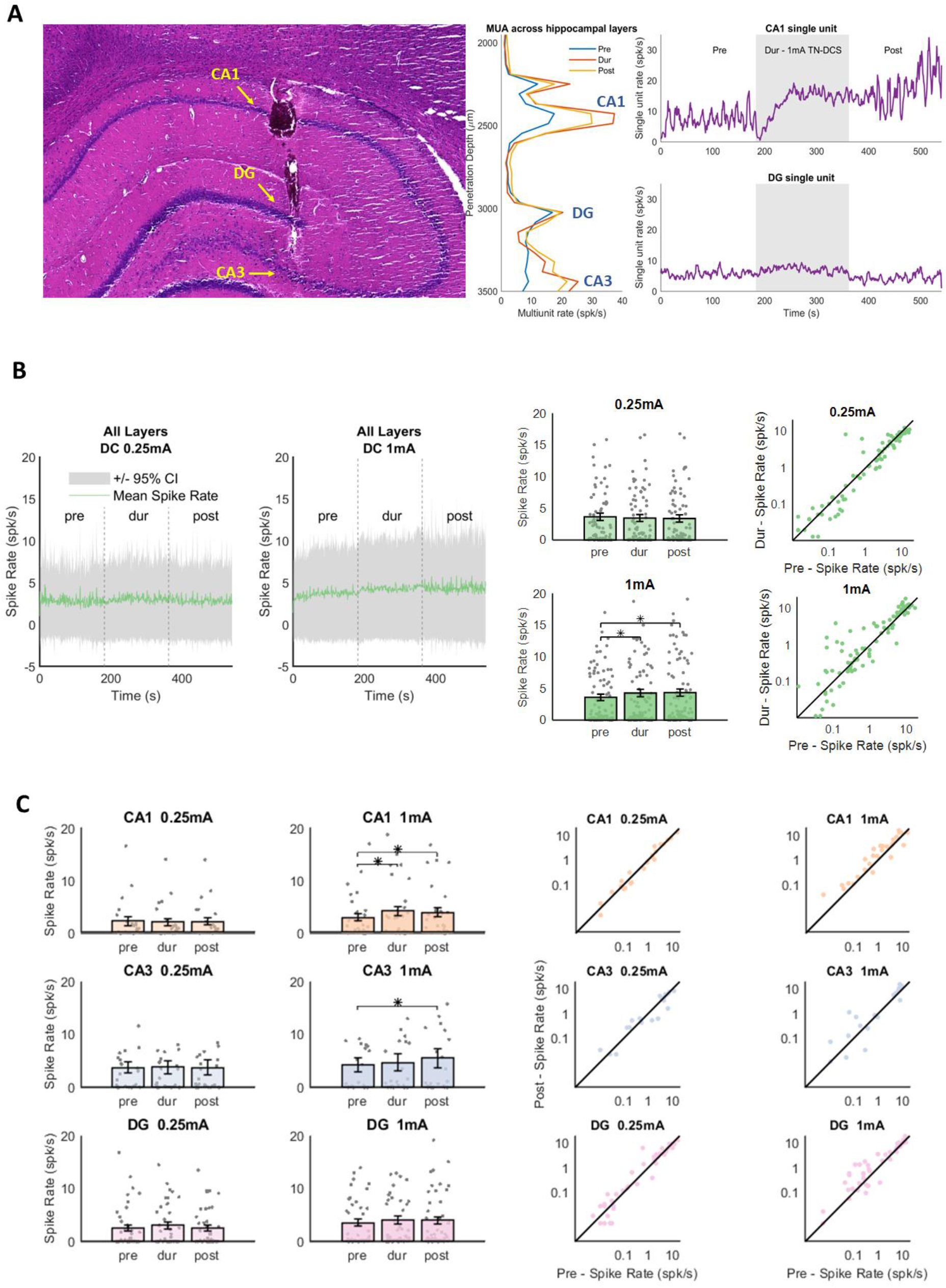
Immediate effects of trigeminal nerve direct current stimulation effects on hippocampal spike-rate. A. Left panel shows histological verification of electrode track traversing different hippocampal layers. Middle panel shows multi-unit activity (MUA) across hippocampal layers in the pre (blue), during (red) and post (yellow) conditions. The peaks in MUA correspond with the CA1, CA3 and DG layers visible in the histology. Right panel shows example single-unit recordings from a putative CA1 and DG neuron showing spike-rate across time. The CA1 neuron spike-rate increases during 1 mA TN-DCS (grey box) and continues to rise after TN-DCS is switched off. In contrast, spike-rate of the DG neuron showed little to no change during TN-DCS. B. Spike-rate for all neurons (CA1, CA3 and DG) during 0.25mA (n=87) and 1mA (n=100) TN-DCS. Left panels show average spike-rate (green lines) with confidence intervals (grey box). Middle panels show mean spike-rate in the pre, during and post conditions. 1 mA TN-DCS caused a significant increase in spike-rate (p values: Pre-Dur: 0.007, Post-Dur: 0.417, Pre-Post: p<0.001), but 0.25 mA TN-DCS did not. The right panels shows the pre versus during spike rate for all neurons. Note for 1 mA TN-DCS there are more points are above the diagonal which is not the case for 0.25 mA TN-DCS. C. A similar analysis for the same neurons in B but now divided into groups based on hippocampal layer (CA1, CA3 and DG). 1mA TN-DCS increase spike rate in CA1 (29 neuron) and CA3 (24 neuron). 0.25mA stimulation had no effect on spike rate in any layer. 1mA TN-DCS caused spike rate in CA1 and CA3 to increase in the during and post condition (CA1 neurons, p values: Pre-Dur: 0.028, Pre-Post: 0.010, Dur-Post: 1.328 ; CA3 neurons, p values: Pre-Dur: 0.874, Pre-Post: 0.039, Dur-Post: 0.215 ). There was no effect of 1 mA TN-DCS on DG neurons. For 1mA, putative CA1 and CA3 layer more points are above the diagonal which is not the case for 1mA TN-DCS in DG layer, 0.25 mA TN-DCS in putative CA1, CA3 and DG layer. Conditions with a significant difference a indicated with an *.

### Electrical stimulation setup

To deliver TN-DCS a rectangular metal electrode (1cm^2^) was coated in gel (Signa Gel, Parker Labs, New Jersey) and attached to the lower jaw to target the marginal branch of trigeminal nerve. The return electrode (1cm^2^) was also coated with gel and attached to the proximal part of the tail. The jaw electrode was connected to the positive terminal of a current source (AM 2200 analog stimulus isolator, A-M Systems, Sequim, WA) and the tail electrode was connected to the negative terminal. The current source was controlled via an analog voltage waveform generated through an output channel on a data acquisition card (NI USB-6216, National Instruments, Austin, TX) at a sampling rate of 100 kHz. The data acquisition card was connected to a standard PC and controlled via custom MATLAB software.

### Experimental protocol

Neural activity was recorded in 9-minute blocks consisting of a 3-minute pre-stimualtion, a 3-minute during and a 3-minute post-stimulation condition. During stimulation we applied DC at either 0.25 mA or 1 mA. To monitor long-term effects, after the 9-minute recording a 3-minute block of neural activity was recorded every 15 minutes for up to 1 hour.

### Blockade experiments

We performed two blockade experiments. The experimental protocol described above was repeated with one difference: In blockade experiment 1 (n = 2 rats), 10 minutes prior to the start of the experimental protocol, we subcutaneous injected xylocaine (1.0ml, 2% solution) to block the same marginal branch of the trigeminal nerve. In blockage experiment 2 (n = 5 rats), 15 minutes prior to the start of the experimental protocol we gave intraperitoneal clonidine (0.05 mg/kg; Sigma-Aldrich, USA) which inhibits LC activity [42]. In both blockage experiments, we stimulated for 3-minutes at 1 mA.

### Histology

After recording, 5 seconds of 30μA DC was passed through the deepest electrode as well as the most superficial electrode. This procedure effectively marked the track of the silicon probe. We then performed a transcardial perfusion (4 % paraformaldehyde VWR Chemicals) where the brain was extracted, fixed in 4% PFA for 24 hours, and embedded in paraffin. We then sliced the brain into 10-μm slices and stained with hematoxylin and eosin.

### Extracting single units

We performed spike sorting using SpykingCircus (https://github.com/spyking-circus/spyking-circus ) [43]. Spyking Circus took in the raw 32-channel recordings and via a series of automated steps clustered neural activity into putative single-unit data. We then performed manual curation using Phy viewer (https://github.com/cortex-lab/phy ). We then extracted spike-times from only well isolated single-units and calculated spike-rates per condition (pre, during and post).

### Spike train thinning

Spike train and LFP data were down sampled to 1000 Hz. The spike-field coherence and spike triggered average metrics are sensitive to spike rate [44]. Therefore, to avoid bias we performed spike thinning [44] by randomly removing spikes until the spike rate of all stimulation conditions (pre, during and post) matched the spike rate of the condition with the lowest spike rate at the beginning of the thinning procedure.

### Spike triggered average analysis

For each single-unit the spike-triggered average (STA) between that single-unit and the LFP on all 32 electrodes, for each stimulation condition (pre, during and post), was calculated as follows,

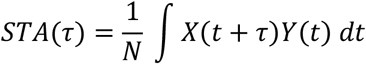

where *X* was the timeseries of LFP on the electrode in question, *Y* was the spike train timeseries represented as a sum of Dirac delta functions, *τ* was the time window within which STA was assessed and was set to ±1000ms, t was time and N was the number of samples. The power spectrum of the STA was then extracted and finally the STA power in the theta (4-8Hz) and gamma (30-80Hz) bands were calculated.

### Spike-field coherence analysis

For each single-unit the spike-field coherence (SFC) between that single-unit and the LFP on all 32 electrodes, for each stimulation condition (pre, during and post), was calculated as follows,

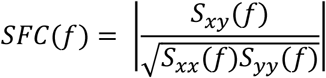

where *S*_*xx*_ was the power spectral density of the LFP on the electrode in question, *S*_*yy*_ was the power spectral density of the spike train, and *S*_*xy*_ was the cross-power spectral density between the sike-train and the LFP. Finally, the coherence in the theta (4-8Hz) and gamma (30-80Hz) bands were calculated.

### Statistics

The effect of TN-DCS amplitude (0.25 mA and 1 mA) and stimulation condition (pre, during and post) on spike-rate was analyzed using a linear mixed model with neuron number and rat as random effects and amplitude and stimulation condition as fixed effects [45], (MATLAB, fitlme.m, model syntax Wilkinson notation: Spike-rate ∼ Amplitude*Condition +Amplitude + Condition + (1|Neuron) + (1|Rat)). All linear mixed model fits were verified by checking the normality of the residuals. Full linear model tables are reported in supplementary. ANOVA was then performed on the model output to test the significance of the fixed effects and their interactions. ANOVA F-statistics and p-values for the fixed effects and interactions are reported under Results. The number of post-hoc test comparisons was 3: pre-during, during-post and pre-post.

The effect of 1mA TN-DCS on spike rate over longer time conditions (pre, dur, post, post15, post30, post45 and post60 mins) was analyzed using a linear mixed model (Spike-rate ∼ conditions + (1|Neuron) + (1|Rat)). ANOVA was then performed on the model output to test the significance of the fixed effect. The number of post-hoc test comparisons was 21 (i.e. all possible comparisons for the 7 different conditions).

To test the effect of 1 mA TN-DCS on a range of synchronization metrics different linear mixed models were used. In each case the stimulation condition (pre, during or post) was set as a fixed effect and neuron number and rat as random effects (Synchronization Metric ∼ Condition + (1|Neuron) + (1|Rat)). The synchronization metric was then set as either: STA power theta band, STA power gamma band, SFC theta band or SFC gamma band. ANOVA was performed on the model output to test the significance of the fixed effects. The number of post-hoc test comparisons was 3: pre-during, during-post and pre-post.

Post-hoc testing was conducted using the two-sided Wilcoxon signed rank. All reported p-values were Bonferroni corrected for multiple comparisons using the number of test comparisons stated above. Alpha of all tests was set to 0.05 and effect sizes are reported as Cohen’s d [46].

## Result

### Effect of TN-DCS on hippocampal spike-rate

Fig. 1A shows an example of how the peaks in the multi-unit activity typically matched with layered structure of the hippocampus. On the basis of this multi-unit activity pattern, combined with recording depth and histology we putatively assigned each single-unit (after spike-sorting) to a specific hippocampal layer. The right panel on Fig. 1A shows the spike-rate over time for two single-units, putatively assigned to DG and CA1. In this example, the two single-units exhibited different spike-rate effects during TN-DCS. In Fig. 1B the left two panels show spike-rate over time averaged across all single-units during 0.25 (n = 87) and 1 mA (n = 100) TN-DCS. The middle panels show the spike-rate in the three conditions (pre, during, post) for each single-unit for 0.25 and 1 mA. An ANOVA of the linear mixed model fit to this data showed that spike-rate was significantly affected by TN-DCS amplitude (FStat= 5.93, p= 0.015). Stimulation condition (pre, during, post) alone was not a significant fixed effect (FStat= 1.12, p= 0.326), but there was a significant interaction between TN-DCS amplitude and stimulation condition (FStat= 3.31, p= 0.037).There were no significant differences between the pre, during and post intercepts. The post condition slope was significantly different from zero, the slopes between post and pre were significantly different and the slopes between post and during were not significantly different (full model details are provided in Supplementary). For the 1 mA TN-DCS data, post-hoc testing (Wilcoxon signed-rank) showed that spike-rate in the during condition was significantly higher than in the pre-condition (z=3.06, p=0.007, Cohen’s d=0.12), post spike-rate was significantly higher than the pre (z=4.61, p<0.001, Cohen’s d=0.13) but during and post spike-rates were not significantly different (z=1.48, p=0.417, Cohen’s d=0.02). The right panels in Fig. 1B show that for 1 mA TN-DCS the during condition (y-axis) tended to have higher spike-rates than the pre-condition. This was not the case for 0.25 mA TN-DCS.

Fig. 1C shows the same analysis but now repeated with the neurons divided into putative groups of CA1 (0.25 mA n=23, 1 mA n=29), CA3 (0.25 mA n=24, 1 mA n=24) and DG (0.25 mA n=40, 1 mA n=47) neurons. For CA1 neurons, ANOVA of the linear mixed model showed that spike-rate was significantly affected by TN-DCS amplitude (FStat= 6.86, p= 0.001). Stimulation condition alone was not significant (FStat= 1.91, p= 0.307), but there was a significant interaction between TN-DCS amplitude and stimulation condition (FStat= 3.13, p= 0.048). For CA3 neurons, ANOVA of the linear mixed model showed that spike-rate was significantly affected by TN-DCS amplitude (FStat= 5.88, p= 0.016) but neither stimulation condition (FStat= 0.12, p= 0.879) nor the interaction between TN-DCS amplitude and stimulation condition (FStat= 1.23, p= 0297) were significant. For DG neurons none of the fixed effects were significant (amplitude FStat= 0.25, p= 0.616; condition FStat= 0.26, p= 0.772; interaction FStat= 0.39, p= 0.675). Post-hoc testing (Wilcoxon signed-rank) for the 1 mA CA1 and the 1 mA CA3 data was performed. For CA1 neurons at 1 mA TN-DCS, spike-rate in the during condition was significantly higher than in the pre-condition (z=3.17, p=0.028, Cohen’s d=0.24); post spike-rate was significantly higher than the pre (z=2.99, p=0.010, Cohen’s d=0.19) but the during spike-rate was not significantly different from the post (CA1 z=0.76, p=1.328, Cohen’s d=0.05). For CA3 neurons at 1 mA TN-DCS post spike-rate was significantly higher than the pre (z=2.48, p=0.039, Cohen’s d=0.15). However, spike-rate in the during condition was not different from the pre (z=1.05, p=0.874, Cohen’s d=0.05) nor post conditions (CA1 z=1.80, p=0.215, Cohen’s d=0.09).

### Xylocaine and clonidine blockage experiments

To test whether the observed effects were directly caused by TN stimulation or indirectly through volume conduction of the DC stimulation to the hippocampus we used subcutaneous injections of xylocaine to block the TN. We tested whether TN-DCS effects were caused by TN or through volume conduction by blocking the nerve using xylocaine. We then applied the same 1mA TN-DCS protocol. Results are shown in Fig. 2, left column (n=2 rats, n=39 neurons). ANOVA of the linear mixed model fit (full results in Supplementary) showed no effect of stimulation condition on spike-rate after TN blockage (FStat=0.18, p=0.834), indicating that TN-DCS effects on hippocampal spike-rate (shown in Fig. 1) can be directly attributed to TN activation.

**Figure 2.**
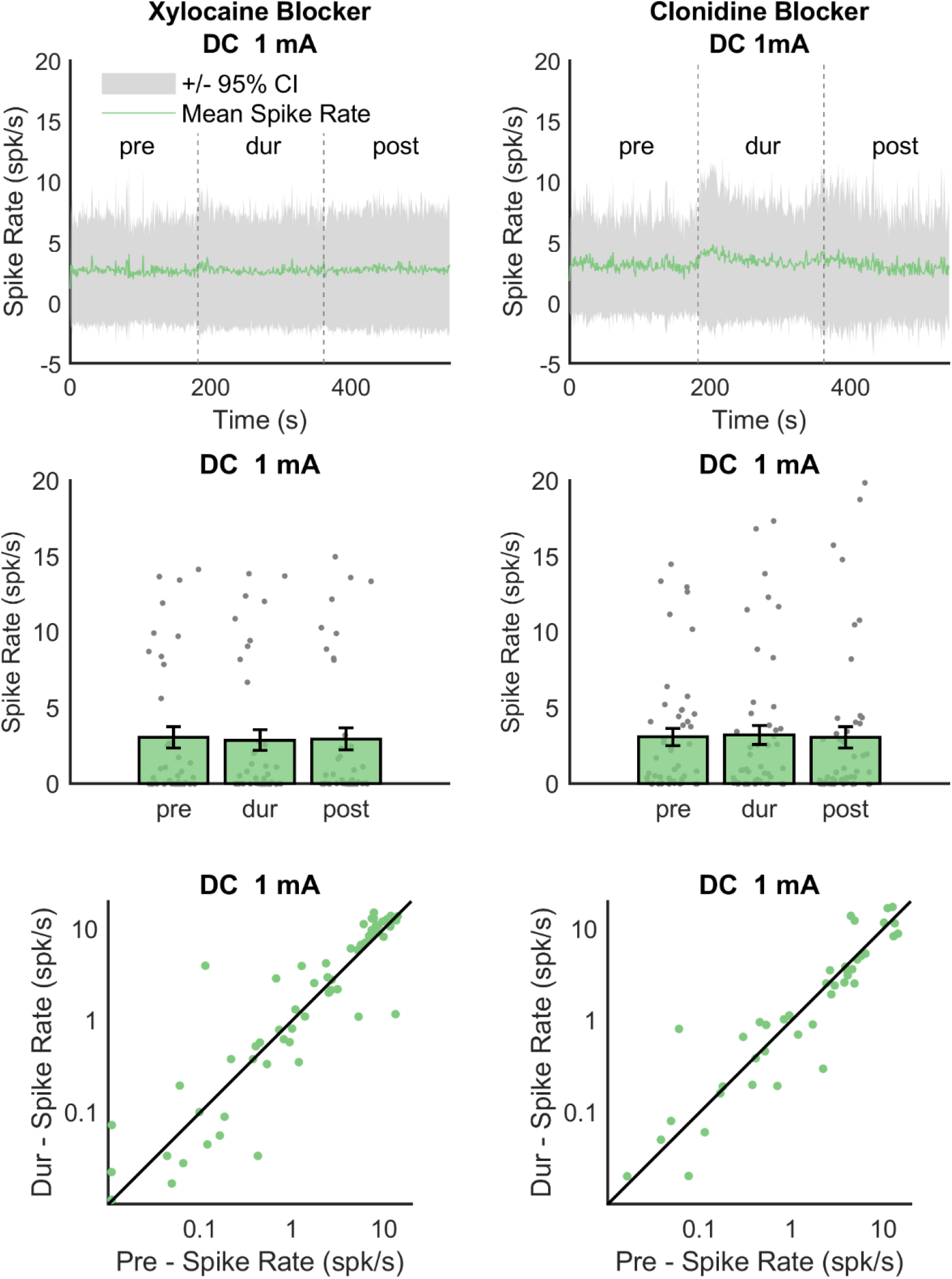
Xylocaine and clonidine blockage experiments. Xylocaine: After trigeminal nerve blockage 1 mA TN-DCS has no effect on hippocampal spike-rate. Upper panel shows average spike-rate over time (green line) and 95% confidence intervals (grey). Middle panel shows mean spike rate in the pre, during and post conditions. There was no significant effect of condition. Bottom panel shows pre versus during spike rate for all neurons. Clonidine: After clonidine injection1 mA TN-DCS has no effect on hippocampal spike-rate. Same convention as xylocaine data. Again, no significant effect of condition was observed.

In a separate experiment we tested whether TN-DCS effects were mediated through the LC-noradrenergic system by injecting clonidine to suppress LC activity [47, 48]. The results are shown in Fig. 2, right column (n=5 rats, n=48 neurons). ANOVA of the linear mixed model fit (full results in Supplementary) showed no effect of stimulation condition on spike-rate after clonidine injection (FStat=0.10, p=0.904), indicating that TN-DCS effects on hippocampal spike-rate (shown in Fig. 1) may be caused by an LC-noradrenergic pathway activation.

### TN-DCS causes prolonged increase in hippocampal spike rate

Hippocampal spike-rate increases outlasted the during-stimulation condition. Therefore, in a follow-up experiment we tracked spike-rate over a period of 1 hour. Fig. 3 shows the spike-rate for data recorded in n=88 neurons, n=5 rats. ANOVA of the linear mixed model fit (full results in Supplementary) showed a significant effect of condition (FStat=3.16, p=0.004). Post-hoc testing (Wilcoxon signed-rank) showed that spike-rates between the following conditions were significantly different: pre and post (z=4.27, p<0.001, Cohen’s d=0.11); pre and post15 (z=3.97, p=0.002, Cohen’s d=0.19); dur and post15 (z=3.27, p=0.023, Cohen’s d=0.12); post15 and post45 (z=-3.53, p=0.009, Cohen’s d=-0.17); post15 and post60 (-z=3.68, p=0.005, Cohen’s d=-0.28); post30 and post60 (z=-3.34, p=0.017, Cohen’s d=-0.22); post45 and post60 (z=-3.22, p=0.027, Cohen’s d=-0.12). All other comparisons were not significantly different.

**Figure 3.**
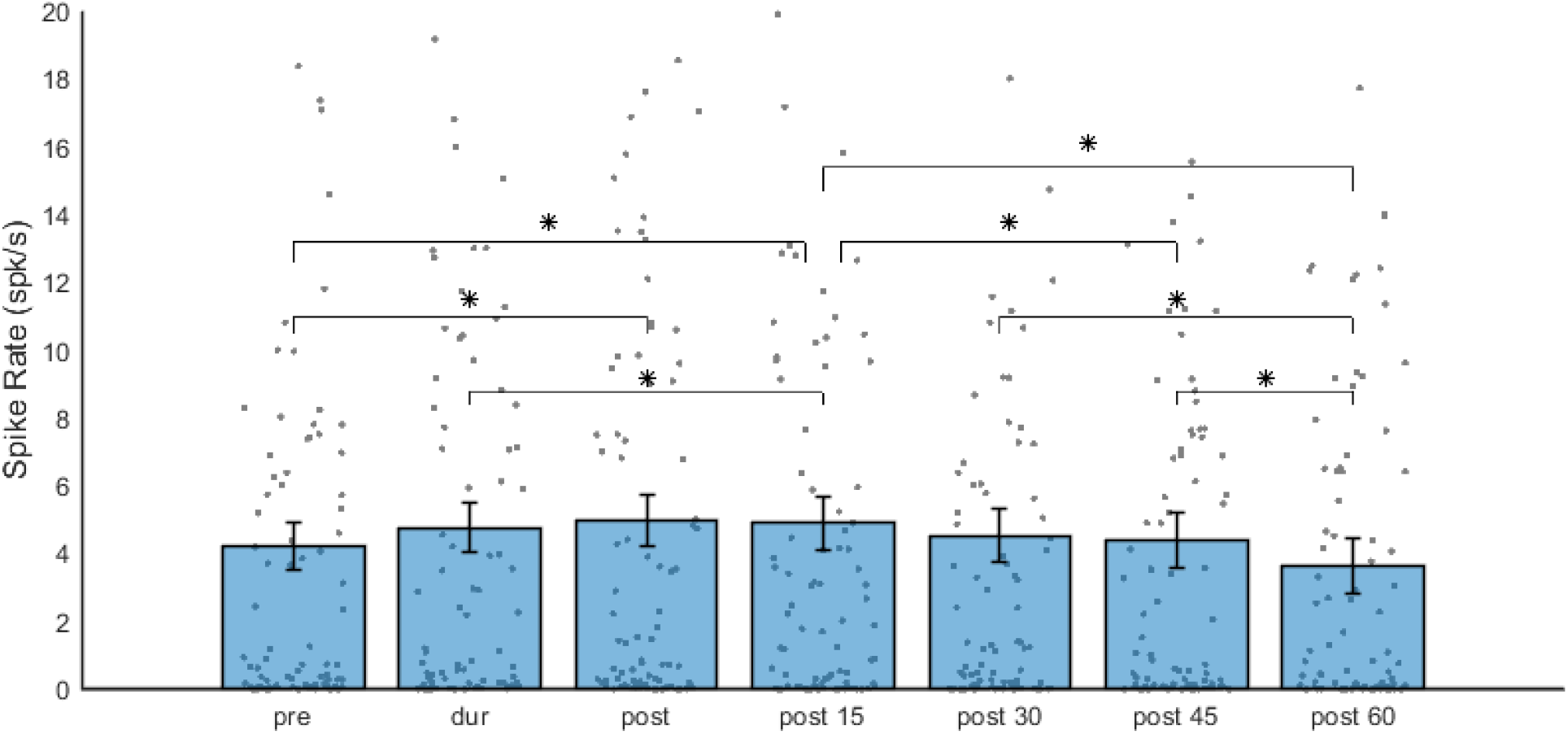
Long-term effects of trigeminal nerve direct current stimulation on hippocampal spike-rate. 1 mA TN-DCS caused a sustained increase in spike-rate lasting up to 45 minutes (88 neurons) before returning to spike-rates similar to the pre-condition. Conditions with a significant difference a indicated with an *.

### TN-DCS affects both STA and SFC

Fig. 4A shows the STA and SFC for an individual putative CA3 neuron at -3500 μm depth. The two left most panels shows the STA and the SFC computed between this neuron and the LFP recorded on the same electrode (i.e. LFP depth was also -3500 μm). The two right most panels of Fig.4A shows the STA and SFC computed between this neuron and the LFP at a different depth (in this case -2950 μm). STA and SFC for the pre (blue), during (red) and post (yellow) conditions are shown. In this neuron 1 mA TN-DCS causes an increase in the STA which remains present after TN-DCS was switched off. Similarly, TN-DCS appears to cause an increase the SFC in the theta and gamma bands. For each individual neuron, we computed the STA and SFC using the LFPs recorded on all 32 electrodes spanning the different hippocampal layers. Fig. 4B shows the SFC (coherence encoded as color, frequency x-axis) across all 32 electrodes (y-axis) for the same neuron as in panel A. Here, we observe that the coherence between this neuron (location indicated by the grey line) and the LFP is higher on some electrodes and lower on others.

**Figure 4.**
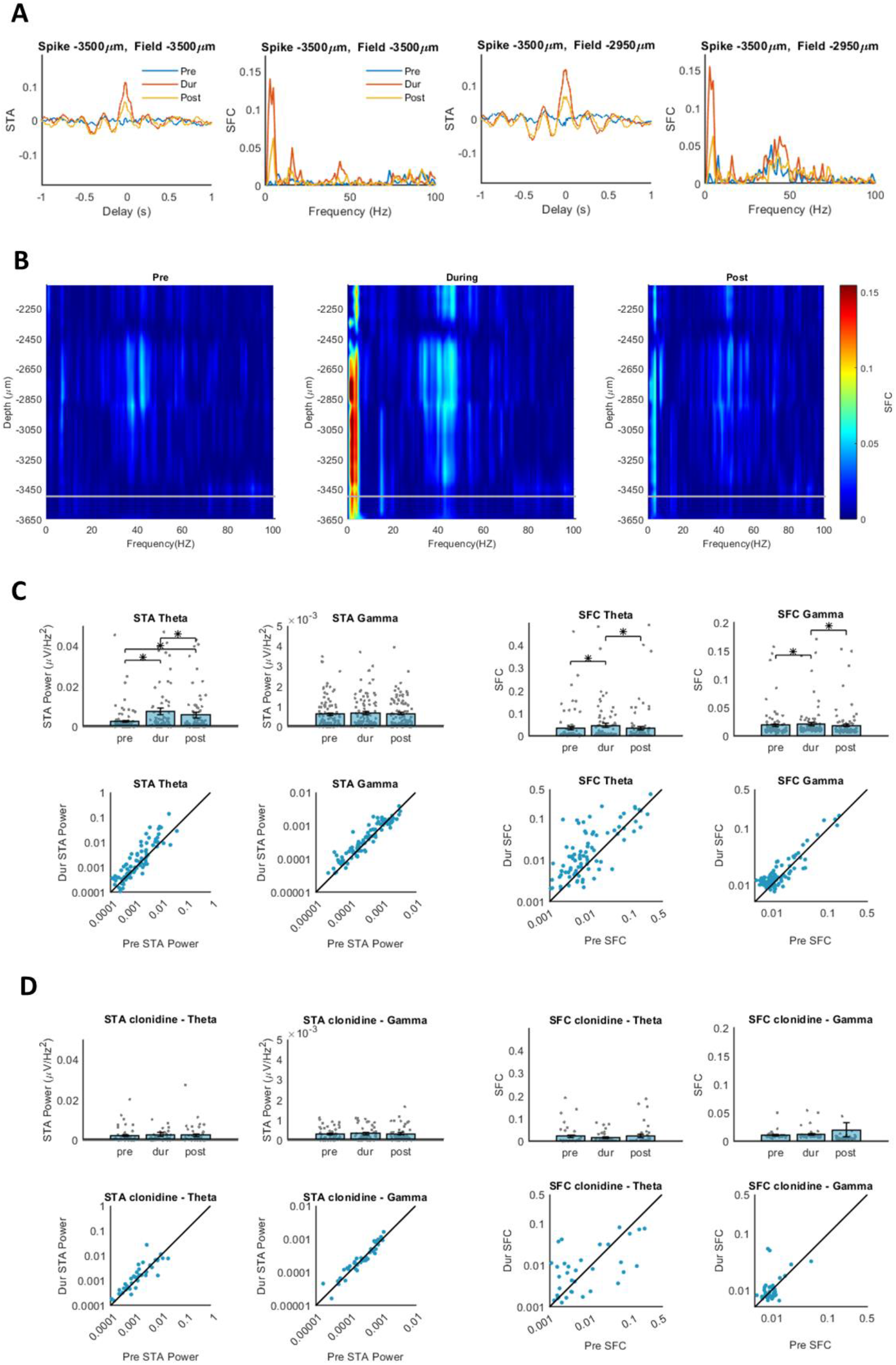
Trigeminal nerve direct current stimulation effects on spike-triggered average and spike-field coherence. A. Individual example of STA and SFC for one single neuron recorded from CA3 (at -3500 μm depth). The two left most panels show STA and SFC calculated using the LFP recorded on the same channel as the neuron, while the two right most panels show STA and SFC calculated using the LFP recorded on a different channel. In this example 1 mA TN-DCS causes an increase in both the STA and SFC in the during (red) condition compared to the pre (blue). The effect persists after TN-DCS is switched off (yellow). B. Individual example of spike-field coherence for one single neuron across LFPs on all 32 electrodes. During stimulation coherence between a CA3 neuron (grey horizontal line) and LFPs on some channels increased among theta to delta band and gamma band (middle panel). After stimulation (right panel), this coherence decreased. Notice how coherence between this CA1 neuron and DG fields was largest before and after stimulation and extended into the theta range. C. At the group level (100 neurons) spike trigger average (STA) in theta band increased during stimulation as compared to the pre-condition (post-hoc z=7.35, p<0.001, Cohen’s d=0.40), after stimulation STA decreased as compare to the during condition (during-post z=3.37, p<0.002, Cohen’s d=0.10 ) but was still higher than the pre-condition (pre-post z=3.52 p=0.001, Cohen’s d=0.26) . At the group level (91 neuron) spike-field coherence (SFC) increased in the theta band (post-hoc pre-during z=4.43, p<0.001, Cohen’s d=0.13) and in the gamma band (post-hoc pre-during z=3.38, p<0.001, Cohen’s d=0.01). SFC theta and SFC gamma then decreased in the post condition (theta during-post z=4.23, p<0.001, Cohen’s d=0.16, gamma during-post z=4.66, p<0.001, Cohen’s d=0.10). D. STA and SFC in clonidine blockade experiment (n=48 neurons for STA, n=39 neurons for SFC). After injection of clonidine there was no effects of 1mA TN-DCS on either STA theta power (p=0.80) or STA gamma power (p=0.19), nor did we observe an effect of 1mA TN-DCS on either SFC in the theta band (p=0.18) or the SFC in the gamma band (p=0.63).

Because the gamma and theta bands are critical in hippocampal memory encoding [33, 35, 36] we focused on the STA and SFC in these two bands. For each neuron, we picked the neuron-electrode pair that showed the highest value for each of the four metrics (either STA theta, STA gamma, SFC theta or SFC gamma) in the during condition for further analysis. For this neuron-electrode pair, we then quantified differences in the pre, during and post conditions using linear mixed models. We then performed post-hoc testing (Wilcoxon signed-rank) when a significant main-effect was detected. The results are shown in Fig 4C (n=6 rats, n=100 neuron-electrode pairs for STA, n=91 neuron-electrode pairs for SFC). We found that STA theta power significantly increased during 1 mA TN-DCS (for full linear model results see Supplementary - FStat=9.40, p<0.001; post-hoc pre-during z=7.35, p<0.001, Cohen’s d=0.40), it then decreased in the post condition (during-post z=3.37, p<0.002, Cohen’s d=0.10) but still remained higher than in the pre condition (pre-post z=3.52 p=0.001, Cohen’s d=0.26). However, no significant effect of 1 mA TN-DCS on STA gamma power was found (FStat=0.84, p<0.434). We found that TN-DCS caused a significant increase SFC in the theta band (FStat=6.67, p=0.001; post-hoc pre-during z=4.43, p<0.001, Cohen’s d=0.13) and on SFC in the gamma band (FStat=6.28, p<0.002; post-hoc pre-during z=3.38, p<0.001, Cohen’s d=0.01). SFC theta and SFC gamma then decreased in the post condition (theta during-post z=4.23, p<0.001, Cohen’s d=0.16, gamma during-post z=4.66, p<0.001, Cohen’s d=0.10).

To investigate a potential role of the LC-noradrenergic pathway in mediating the observed effects of TN-DCS on STA and SFC we now repeated this experiment and analysis after injection of clonidine. The results are shown in Fig. 4D (n=5 rats, n=48 neuron-electrode pairs for STA, n=39 neuron-electrode pairs for SFC). After injection of clonidine we found no effects of 1mA TN-DCS on either STA theta power (FStat=0.22, p=0.80) or STA gamma power (FStat=1.68,p=0.19), nor did we observe an effect of 1mA TN-DCS on either SFC in the theta band (FStat=1.75,p=0.18) or the SFC in the gamma band (FStat=0.47,p=0.63).

## Discussion

In a standard tDCS montage the supraorbital electrode will pass current through the skin on the forehead, thus stimulating the trigeminal nerve. We hypothesized that 1) DC stimulation of the trigeminal nerve could modulate hippocampal activity and 2) this modulation occurs via an LC-noradrenergic pathway. In this study, we tested these hypotheses using a rat model. Firstly, we showed that there is a functional pathway from the trigeminal nerve to hippocampus which can modulate neural activity. Secondly, we used a blocker experiment to show this modulation appears to depend on the LC-noradrenergic pathway. More specifically, we found that 1mA TN-DCS caused significant increases in hippocampal spike-rate while 0.25 mA TN-DCS did not. Moreover, the effect of 1mA TN-DCS on hippocampal spike-rate persisted for up to 45 minutes. Blocking the TN via xylocaine injection prevented the hippocampal response to TN-DCS, indicating the observed effects were directly caused by TN stimulation. Blocking the LC-noradrenergic pathway via i.p. injection of clonidine [42] also prevented the response, indicating the LC-noradrenergic pathway may play a central role in mediating this response. Additionally, TN-DCS increased hippocampal SFC in both the theta and gamma bands and STA power in the theta band. These results support the hypothesis that TN-DCS can modulate hippocampal neural circuits that underly learning and memory processes.

tDCS subjects often experience a tingling or itching sensation due to the involvement of the thalamic pathway to the sensory cortex [21]. This sensation arises from the activation of sensory fibers in the scalp’s trigeminal and occipital nerves [15] indicating that cranial and cervical nerves are activated during tDCS. However, the potential neuromodulatory role of this bottom-up pathway is only now beginning to be studied. Vanneste et al. [17, 31, 32] linked tDCS effects on memory to stimulation of the greater occipital nerve thereby pointing to the LC-noradrenergic pathway. We aimed to further explore this bottom-up pathway and have recently shown TN-DCS directly activates the trigeminal nuclei [49]. These nuclei project to several other brainstem nuclei including the LC which plays a central role in controlling noradrenaline release in several brain regions, including the hippocampus [23]. In this study we further elucidate this bottom-up pathway by being the first group to show that TN-DCS directly modulates neural activity in the hippocampus. Indeed, our results showed that TN-DCS leads to increased spike-rates in the hippocampus, in addition to an increase in SFC in frequency bands which are relevant for hippocampal memory formation [33].

### Xylocaine blockade experiments

We tested whether TN-DCS effects could have been mediated through volume conduction to the hippocampus by inactivating the trigeminal nerve using xylocaine [49, 50]. After TN blockade, we observed no hippocampal response to 1mA TN-DCS. Thus, these findings support the hypothesis that the observed effects in the hippocampus were caused by TN stimulation rather than volume conduction.

### Clonidine blockade experiments

We tested the hypothesis that the effects of TN-DCS on hippocampal activity are mediated by the LC-noradrenergic pathway by injecting the α-2 adrenergic agonist clonidine which specifically blocks LC activity [42, 48]. We observed no significant changes in spike-rate nor in SFC during 1 mA TN-DCS. Thus, our results support a role for the LC-noradrenergic pathway as a viable player in the TN-DCS bottom-up neuromodulation pathway. However, our experiments did not completely rule out the involvement of other potential pathways such as the nucleus of the solitary tract, raphe nuclei, and their projections to hippocampus or thalamic and somatosensory cortex input to CA1 [21, 23]. We also cannot rule out the potential inhibitory effects of clonidine working directly on α-2 adrenergic receptors in the hippocampus. However, to the best of our knowledge, there is currently no evidence that clonidine directly inhibits hippocampal activity. Indeed, our own data show that hippocampal spike-rate remains similar even after injection of clonidine (pre spike-rate in Fig. 1B and Fig. 2B are similar). On the other hand many studies have shown that clonidine and other α-2 adrenergic receptor antagonists directly suppress LC activity [51, 52]. Finally, we believe more experiments are needed to fully understand this complex pathway.

### STA and SFC analysis

In our STA and SFC analysis we focused on the theta and gamma bands as they play a central role in hippocampal memory formation [33, 36, 53]. Our results showed that both STA and SFC in the theta band increased during TN-DCS stimulation and decreased after stimulation. Theta oscillations play a significant role in cognitive processes, particularly in the encoding and retrieval of episodic and spatial memories [54]. Additionally, we also observed an increase in SFC gamma band during TN-DCS, which returned to baseline after stimulation was switched off. Gamma oscillations in the hippocampus also play a significant role in driving spike-timing-dependent plasticity and for the formation of coherent and integrated memories [53, 55]. However, any effect of TN-DCS on gamma band in the STA did not reach significance. This may be because the STA is an averaging method dominated by the largest amplitude fluctuations in the LFP, and theta oscillations have a larger amplitude than gamma because of 1/f LFP scaling. The SFC analysis is conducted in the frequency domain meaning it meaning it may be less dominated by larger amplitude, lower frequency LFPs. Interestingly, no effects of TN-DCS on the STA and SFC were present when a clonidine blocker was used, again suggesting that LC-noradrenergic projections could play a key role in controlling important hippocampal memory circuits [27, 56].

### TN-DCS effect sizes

Most studies using tDCS on human participants tend to report small effect sizes [57–59]. Interestingly, we observed that the Cohen’s d effect sizes of TN-DCS on hippocampal activity ranged between 0.19 and 0.53, with the mean being 0.26. These are considers to be small effect sizes [60]. Thus, the effect sizes expected from the TN-DCS bottom-up pathway found in our study are consistent with the typical effect sizes observed in human tDCS studies.

## Conclusion

Recent evidence suggests that tDCS memory effects may be caused via a bottom-up pathway in which nerves in the scalp are stimulated [17, 61] and not via the electric field in the brain as is generally assumed. Our results show that DC stimulation of the TN does affect hippocampal activity in a rat model and thus provide a neural basis to support the bottom-up pathway. Thus, TN-DCS emerges as a potent tool for memory manipulation which could hold promise for cognitive interventions. However, it is important to note that our work focused on illuminating the tDCS bottom-up pathway and did not rule out a role for the electric field in the brain in humans tDCS studies. Further studies will be needed to fully disentangle both of these tDCS mechanisms.

## Acknowledgements

This work was supported by FWO Funding G0B4520N and NIH Funding 1R01MH123508-01. Liyi Chen was fund by China Scholarship Council at CSC202006380043, Zhengdao Deng was funded by PhD fellowship at EFD-LSMDT5-02010 AND QPG-327008-NUTTIN-FWO-SBO-SCATMAN.

## Conflict of interest declaration

All authors confirm there are no conflicts of interest that could have influenced the outcome of this work.

## Notes

### Competing Interest Statement

The authors have declared no competing interest.

## Reference

1. Nitsche MA, Paulus W. Excitability changes induced in the human motor cortex by weak transcranial direct current stimulation. J Physiol. 2000;527:633.

2. Horvath JC, Forte JD, Carter O. Quantitative Review Finds No Evidence of Cognitive Effects in Healthy Populations From Single-session Transcranial Direct Current Stimulation (tDCS). Brain Stimul. 2015;8:535–550.

3. Lefaucheur JP, Antal A, Ayache SS, Benninger DH, Brunelin J, Cogiamanian F, et al. Evidence-based guidelines on the therapeutic use of transcranial direct current stimulation (tDCS). Clin Neurophysiol. 2017;128:56–92.

4. Buch ER, Santarnecchi E, Antal A, Born J, Celnik PA, Classen J, et al. Effects of tDCS on motor learning and memory formation: A consensus and critical position paper. Clin Neurophysiol. 2017;128:589–603.

5. Hill AT, Fitzgerald PB, Hoy KE. Effects of Anodal Transcranial Direct Current Stimulation on Working Memory: A Systematic Review and Meta-Analysis of Findings From Healthy and Neuropsychiatric Populations. Brain Stimul. 2016;9:197–208.

6. Javadi AH, Cheng P. Transcranial Direct Current Stimulation (tDCS) Enhances Reconsolidation of Long-Term Memory. Brain Stimul. 2013;6:668–674.

7. Ferrucci R, Mameli F, Guidi I, Mrakic-Sposta S, Vergari M, Marceglia S, et al. Transcranial direct current stimulation improves recognition memory in Alzheimer disease. Neurology. 2008;71:493–498.

8. Majdi A, van Boekholdt L, Sadigh-Eteghad S, Mc Laughlin M. A systematic review and meta-analysis of transcranial direct-current stimulation effects on cognitive function in patients with Alzheimer’s disease. Mol Psychiatry 2022 274. 2022;27:2000–2009.

9. Huang Y, Liu AA, Lafon B, Friedman D, Dayan M, Wang X, et al. Measurements and models of electric fields in the in vivo human brain during transcranial electric stimulation. Elife. 2017;6.

10. Louviot S, Tyvaert L, Maillard LG, Colnat-Coulbois S, Dmochowski J, Koessler L. Transcranial Electrical Stimulation generates electric fields in deep human brain structures. Brain Stimul. 2022;15:1–12.

11. Vöröslakos M, Takeuchi Y, Brinyiczki K, Zombori T, Oliva A, Fernández-Ruiz A, et al. Direct effects of transcranial electric stimulation on brain circuits in rats and humans. Nat Commun. 2018;9.

12. Rahman A, Reato D, Arlotti M, Gasca F, Datta A, Parra LC, et al. Cellular effects of acute direct current stimulation: somatic and synaptic terminal effects. J Physiol. 2013;591:2563–2578.

13. Polanía R, Nitsche MA, Paulus W. Modulating functional connectivity patterns and topological functional organization of the human brain with transcranial direct current stimulation. Hum Brain Mapp. 2011;32:1236–1249.

14. Lefaucheur JP, Wendling F. Mechanisms of action of tDCS: A brief and practical overview. Neurophysiol Clin. 2019;49:269–275.

15. van Boekholdt L, Kerstens S, Khatoun A, Asamoah B, Mc Laughlin M. tDCS peripheral nerve stimulation: a neglected mode of action? Mol Psychiatry 2020 262. 2020;26:456–461.

16. Luckey AM, Adcock K, Vanneste S. Peripheral nerve stimulation: A neuromodulation-based approach. Neurosci Biobehav Rev. 2023;149:105180.

17. Vanneste S, Mohan A, Yoo H Bin, Huang Y, Luckey AM, Lauren McLeod S, et al. The peripheral effect of direct current stimulation on brain circuits involving memory. Sci Adv. 2020;6.

18. Khatoun A, Asamoah B, Laughlin MM. Investigating the Feasibility of Epicranial Cortical Stimulation Using Concentric-Ring Electrodes: A Novel Minimally Invasive Neuromodulation Method. Front Neurosci. 2019;13.

19. Asamoah B, Khatoun A, Laughlin MM. ARTICLE tACS motor system effects can be caused by transcutaneous stimulation of peripheral nerves. 10.1038/s41467-018-08183-w.

20. M So PP, Member S, Stuchly MA, Nyenhuis JA. Peripheral Nerve Stimulation by Gradient Switching Fields in Magnetic Resonance Imaging. IEEE Trans Biomed Eng. 2004;51.

21. Stöhr M, Petruch F. Somatosensory evoked potentials following stimulation of the trigeminal nerve in man. J Neurol. 1979;220:95–98.

22. Brunoni AR, Amadera J, Berbel B, Volz MS, Rizzerio BG, Fregni F. A systematic review on reporting and assessment of adverse effects associated with transcranial direct current stimulation. Int J Neuropsychopharmacol. 2011;14:1133–1145.

23. Tyler WJ, Boasso AM, Mortimore HM, Silva RS, Charlesworth JD, Marlin MA, et al. Transdermal neuromodulation of noradrenergic activity suppresses psychophysiological and biochemical stress responses in humans. Sci Rep. 2015;5.

24. Wagatsuma A, Okuyama T, Sun C, Smith LM, Abe K, Tonegawa S. Locus coeruleus input to hippocampal CA3 drives single-trial learning of a novel context. Proc Natl Acad Sci U S A. 2017;115:E310–E316.

25. Kaufman AM, Geiller T, Losonczy A. A Role for the Locus Coeruleus in Hippocampal CA1 Place Cell Reorganization during Spatial Reward Learning. Neuron. 2020;105:1018–1026.e4.

26. Lim EP, Tan CH, Jay TM, Dawe GS. Locus coeruleus stimulation and noradrenergic modulation of hippocampo-prefrontal cortex long-term potentiation. Int J Neuropsychopharmacol. 2010;13:1219–1231.

27. Sara SJ. The locus coeruleus and noradrenergic modulation of cognition. Nat Rev Neurosci. 2009;10:211–223.

28. Kempadoo KA, Mosharov E V., Choi SJ, Sulzer D, Kandel ER. Dopamine release from the locus coeruleus to the dorsal hippocampus promotes spatial learning and memory. Proc Natl Acad Sci U S A. 2016;113:14835–14840.

29. McNamara CG, Dupret D. Two sources of dopamine for the hippocampus. Trends Neurosci. 2017;40:383–384.

30. Ranjbar-Slamloo Y, Fazlali Z. Dopamine and Noradrenaline in the Brain; Overlapping or Dissociate Functions? Front Mol Neurosci. 2020;12.

31. Luckey AM, McLeod SL, Robertson IH, To WT, Vanneste S. Greater Occipital Nerve Stimulation Boosts Associative Memory in Older Individuals: A Randomized Trial. Neurorehabil Neural Repair. 2020;34:1020–1029.

32. Luckey AM, McLeod LS, Huang Y, Mohan A, Vanneste S. Making memories last using the peripheral effect of direct current stimulation. Elife. 2023;12.

33. Lisman JE, Jensen O. The Theta-Gamma Neural Code. Neuron. 2013;77:1002–1016.

34. Fries P. A mechanism for cognitive dynamics: neuronal communication through neuronal coherence. Trends Cogn Sci. 2005;9:474–480.

35. Buzsáki G, Schomburg EW. What does gamma coherence tell us about inter-regional neural communication? Nat Neurosci. 2015;18:484–489.

36. Mysin I, Shubina L. From mechanisms to functions: The role of theta and gamma coherence in the intrahippocampal circuits. Hippocampus. 2022;32:342–358.

37. Lowet E, Sheehan DJ, Chialva U, De Oliveira Pena R, Mount RA, Xiao S, et al. Theta and gamma rhythmic coding through two spike output modes in the hippocampus during spatial navigation. Cell Rep. 2023;42:112906.

38. Benchenane K, Peyrache A, Khamassi M, Tierney PL, Gioanni Y, Battaglia FP, et al. Coherent Theta Oscillations and Reorganization of Spike Timing in the Hippocampal-Prefrontal Network upon Learning. Neuron. 2010;66:921–936.

39. Bacon TJ, Pickering AE, Mellor JR. Noradrenaline Release from Locus Coeruleus Terminals in the Hippocampus Enhances Excitation-Spike Coupling in CA1 Pyramidal Neurons Via β-Adrenoceptors. Cereb Cortex. 2020;30:6135–6151.

40. Hajos M, Hoffmann WE, Robinson DD, Yu JH, Hajós-Korcsok É. Norepinephrine but not serotonin reuptake inhibitors enhance theta and gamma activity of the septo-hippocampal system. Neuropsychopharmacology. 2003;28:857–864.

41. Paxinos G, Watson C. The rat brain in stereotaxic coordinates.(6 th Edtn). Amsterdam, Bost. 2007. 2007.

42. McCune SK, Voigt MM, Hill JM. Expression of multiple alpha adrenergic receptor subtype messenger RNAs in the adult rat brain. Neuroscience. 1993;57:143–151.

43. Yger P, Spampinato GLB, Esposito E, Lefebvre B, Deny S, Gardella C, et al. A spike sorting toolbox for up to thousands of electrodes validated with ground truth recordings in vitro and in vivo. Elife. 2018;7.

44. Lepage KQ, Gregoriou GG, Kramer MA, Aoi M, Gotts SJ, Eden UT, et al. A procedure for testing across-condition rhythmic spike-field association change. J Neurosci Methods. 2013;213:43–62.

45. Yu Z, Guindani M, Grieco SF, Chen L, Holmes TC, Xu X. Beyond t test and ANOVA: applications of mixed-effects models for more rigorous statistical analysis in neuroscience research. Neuron. 2022;110:21–35.

46. Cohen J. Statistical Power Analysis for the Behavioral Sciences. Routledge; 1998.

47. Berridge CW, Page ME, Valentino RJ, Foote SL. Effects of locus coeruleus inactivation on electroencephalographic activity in neocortex and hippocampus. Neuroscience. 1993;55:381–393.

48. Totah NK, Neves RM, Panzeri S, Logothetis NK, Eschenko O. The Locus Coeruleus Is a Complex and Differentiated Neuromodulatory System. Neuron. 2018;99:1055–1068.e6.

49. Majdi A, Asamoah B, Laughlin MM. Understanding Neuromodulation Pathways in tDCS: Brain Stem Recordings in Rat During Trigeminal Nerve Direct Current Stimulation. BioRxiv. 2023:2023.09.14.557723.

50. Kerstens S, Orban de Xivry JJ, Mc Laughlin M. A novel tDCS control condition using optimized anesthetic gel to block peripheral nerve input. Front Neurol. 2022;13.

51. Berridge CW, Foote SL. Effects of locus coeruleus activation on electroencephalographic activity in neocortex and hippocampus. J Neurosci. 1991;11:3135–3145.

52. Yavich L, Jäkälä P, Tanila H. Noradrenaline overflow in mouse dentate gyrus following locus coeruleus and natural stimulation: real-time monitoring by in vivo voltammetry. J Neurochem. 2005;95:641–650.

53. Nyhus E, Curran T. Functional Role of Gamma and Theta Oscillations in Episodic Memory. Neurosci Biobehav Rev. 2010;34:1023.

54. Buzsáki G. Theta Oscillations in the Hippocampus. Neuron. 2002;33:325–340.

55. Fernández-Ruiz A, Oliva A, Nagy GA, Maurer AP, Berényi A, Buzsáki G. Entorhinal-CA3 Dual-Input Control of Spike Timing in the Hippocampus by Theta-Gamma Coupling. Neuron. 2017;93:1213–1226.e5.

56. Eschenko O, Mello-Carpes PB, Hansen N. New Insights into the Role of the Locus Coeruleus-Noradrenergic System in Memory and Perception Dysfunction. Neural Plast. 2017;2017.

57. Schroeder PA, Schwippel T, Wolz I, Svaldi J. Meta-analysis of the effects of transcranial direct current stimulation on inhibitory control. Brain Stimul. 2020;13:1159–1167.

58. Figeys M, Zeeman M, Kim ES. Effects of Transcranial Direct Current Stimulation (tDCS) on Cognitive Performance and Cerebral Oxygen Hemodynamics: A Systematic Review. Front Hum Neurosci. 2021;15:623315.

59. Buch ER, Santarnecchi E, Antal A, Born J, Celnik PA, Classen J, et al. Effects of tDCS on motor learning and memory formation: A consensus and critical position paper. Clin Neurophysiol. 2017;128:589–603.

60. Sawilowsky SS. New Effect Size Rules of Thumb. J Mod Appl Stat Methods. 2009;8:26.

61. Luckey AM, McLeod SL, Mohan A, Vanneste S. Potential role for peripheral nerve stimulation on learning and long-term memory: A comparison of alternating and direct current stimulations. Brain Stimul. 2022;15:536–545.

