## Supplementary for "Trigeminal nerve direct current stimulation causes sustained increase in neural activity in the rat hippocampus"

### Linear Mixed Model Fits

#### Fig 1B - LME Model For All Single Units

**Linear mixed-effects model fit by ML**

**Model information:**

| | |
|---|---|
| Number of observations | 561 |
| Fixed effects coefficients | 6 |
| Random effects coefficients | 106 |
| Covariance parameters | 3 |

**Formula:**

SPK ~ 1 + Amp*Condition + (1 | Neuron) + (1 | Rat)

**Model fit statistics:**

| AIC | BIC | LogLikelihood | Deviance |
|---|---|---|---|
| 2795.9 | 2834.8 | -1388.9 | 2777.9 |

**Fixed effects coefficients (95% CIs):**

| Name | Estimate | SE | tStat | DF | pValue | Lower | Upper |
|---|---|---|---|---|---|---|---|
| {'(Intercept)' } | 4.4748 | 0.57858 | 7.7342 | 555 | 4.9304e-14 | 3.3383 | 5.6113 |
| {'Amp' } | 0.62044 | 0.25471 | 2.4359 | 555 | 0.015169 | 0.12013 | 1.1207 |
| {'Condition_dur' } | -0.12272 | 0.25004 | -0.49079 | 555 | 0.62377 | -0.61387 | 0.36843 |
| {'Condition_pre' } | 0.36795 | 0.25004 | 1.4715 | 555 | 0.14172 | -0.1232 | 0.8591 |
| {'Amp:Condition_dur'} | 0.32711 | 0.333 | 0.98231 | 555 | 0.32637 | -0.32698 | 0.9812 |
| {'Amp:Condition_pre'} | -0.84932 | 0.333 | -2.5505 | 555 | 0.011023 | -1.5034 | -0.19523 |

**Random effects covariance parameters (95% CIs):**

Group: Neuron (100 Levels)

| Name1 | Name2 | Type | Estimate | Lower | Upper |
|---|---|---|---|---|---|
| {'(Intercept)'} | {'(Intercept)'} | {'std'} | 5.4227 | 4.7017 | 6.2543 |

Group: Rat (6 Levels)

| Name1 | Name2 | Type | Estimate | Lower | Upper |
|---|---|---|---|---|---|
| {'(Intercept)'} | {'(Intercept)'} | {'std'} | 2.93e-07 | NaN | NaN |

Group: Error

| Name | Estimate | Lower | Upper |
|---|---|---|---|
| {'Res Std'} | 2.0863 | 1.9559 | 2.2254 |

#### Fig. 1C, top row - LME Model For CA1 Single Units

**Linear mixed-effects model fit by ML**

**Model information:**

| | |
|---|---|
| Number of observations | 156 |
| Fixed effects coefficients | 6 |
| Random effects coefficients | 35 |
| Covariance parameters | 3 |

**Formula:**

SPK ~ 1 + Amp*Condition + (1 | Neuron) + (1 | Rat)

**Model fit statistics:**

| AIC | BIC | LogLikelihood | Deviance |
|---|---|---|---|
| 745.14 | 772.59 | -363.57 | 727.14 |

**Fixed effects coefficients (95% CIs):**

| Name | Estimate | SE | tStat | DF | pValue | Lower | Upper |
|---|---|---|---|---|---|---|---|
| {'(Intercept)' } | 3.5342 | 1.1314 | 3.1238 | 150 | 0.002143 | 1.2987 | 5.7697 |
| {'Amp' } | 1.8303 | 0.69865 | 2.6197 | 150 | 0.0097052 | 0.44979 | 3.2107 |
| {'Condition_pre' } | 1.1125 | 0.73456 | 1.5145 | 150 | 0.13201 | -0.33896 | 2.5639 |
| {'Condition_post' } | 0.36703 | 0.73456 | 0.49966 | 150 | 0.61805 | -1.0844 | 1.8185 |
| {'Amp:Condition_pre' } | -2.2911 | 0.96012 | -2.3862 | 150 | 0.01827 | -4.1882 | -0.39395 |
| {'Amp:Condition_post'} | -0.63987 | 0.96012 | -0.66645 | 150 | 0.50615 | -2.537 | 1.2572 |

**Random effects covariance parameters (95% CIs):**

Group: Neuron (29 Levels)

| Name1 | Name2 | Type | Estimate | Lower | Upper |
|---|---|---|---|---|---|
| {'(Intercept)'} | {'(Intercept)'} | {'std'} | 4.0037 | 2.9404 | 5.4515 |

Group: Rat (6 Levels)

| Name1 | Name2 | Type | Estimate | Lower | Upper |
|---|---|---|---|---|---|
| {'(Intercept)'} | {'(Intercept)'} | {'std'} | 1.5005 | 0.26331 | 8.5508 |

Group: Error

| Name | Estimate | Lower | Upper |
|---|---|---|---|
| {'Res Std'} | 1.8236 | 1.6125 | 2.0624 |

### Fig. 1C, middle row - LME Model For CA3 Single Units

**Linear mixed-effects model fit by ML**

**Model information:**

| | |
|---|---|
| Number of observations | 144 |
| Fixed effects coefficients | 6 |
| Random effects coefficients | 30 |
| Covariance parameters | 3 |

**Formula:**

SPK ~ 1 + Amp*Condition + (1 | Neuron) + (1 | Rat)

**Model fit statistics:**

| AIC | BIC | LogLikelihood | Deviance |
|---|---|---|---|
| 721.08 | 747.81 | -351.54 | 703.08 |

**Fixed effects coefficients (95% CIs):**

| Name | Estimate | SE | tStat | DF | pValue | Lower | Upper |
|---|---|---|---|---|---|---|---|
| {'(Intercept)' } | 3.8719 | 1.5365 | 2.52 | 138 | 0.012874 | 0.83385 | 6.91 |
| {'Amp' } | 1.8452 | 0.76107 | 2.4245 | 138 | 0.016625 | 0.34031 | 3.35 |
| {'Condition_pre' } | 0.040921 | 0.76898 | 0.053215 | 138 | 0.95764 | -1.4796 | 1.5614 |
| {'Condition_post' } | -0.31517 | 0.76898 | -0.40985 | 138 | 0.68255 | -1.8357 | 1.2053 |
| {'Amp:Condition_pre' } | -0.4716 | 1.055 | -0.447 | 138 | 0.65558 | -2.5577 | 1.6145 |
| {'Amp:Condition_post'} | 1.1349 | 1.055 | 1.0758 | 138 | 0.28392 | -0.95116 | 3.2211 |

**Random effects covariance parameters (95% CIs):**

Group: Neuron (24 Levels)

| Name1 | Name2 | Type | Estimate | Lower | Upper |
|---|---|---|---|---|---|
| {'(Intercept)'} | {'(Intercept)'} | {'std'} | 7.0013 | 5.2561 | 9.3259 |

Group: Rat (6 Levels)

| Name1 | Name2 | Type | Estimate | Lower | Upper |
|---|---|---|---|---|---|
| {'(Intercept)'} | {'(Intercept)'} | {'std'} | 8.5513e-06 | NaN | NaN |

Group: Error

| Name | Estimate | Lower | Upper |
|---|---|---|---|
| {'Res Std'} | 1.9382 | 1.7079 | 2.1996 |

### Fig. 1C, bottom row - LME Model For DG Single Units

**Linear mixed-effects model fit by ML**

**Model information:**

| | |
|---|---|
| Number of observations | 261 |
| Fixed effects coefficients | 6 |
| Random effects coefficients | 53 |
| Covariance parameters | 3 |

**Formula:**

SPK ~ 1 + Amp*Condition + (1 | Neuron) + (1 | Rat)

**Model fit statistics:**

| AIC | BIC | LogLikelihood | Deviance |
|---|---|---|---|
| 1339.5 | 1371.6 | -660.75 | 1321.5 |

**Fixed effects coefficients (95% CIs):**

| Name | Estimate | SE | tStat | DF | pValue | Lower | Upper |
|---|---|---|---|---|---|---|---|
| {'(Intercept)' } | 4.7812 | 0.85133 | 5.6161 | 255 | 5.0962e-08 | 3.1047 | 6.4577 |
| {'Amp' } | 0.34231 | 0.6815 | 0.50229 | 255 | 0.6159 | -0.99977 | 1.6844 |
| {'Condition_pre' } | -0.052046 | 0.70543 | -0.07378 | 255 | 0.94124 | -1.4412 | 1.3372 |
| {'Condition_post' } | -0.46346 | 0.70543 | -0.657 | 255 | 0.51178 | -1.8527 | 0.92574 |
| {'Amp:Condition_pre' } | -0.45989 | 0.93521 | -0.49175 | 255 | 0.62332 | -2.3016 | 1.3818 |
| {'Amp:Condition_post'} | 0.36766 | 0.93521 | 0.39313 | 255 | 0.69455 | -1.4741 | 2.2094 |

**Random effects covariance parameters (95% CIs):**

Group: Neuron (47 Levels)

| Name1 | Name2 | Type | Estimate | Lower | Upper |
|---|---|---|---|---|---|
| {'(Intercept)'} | {'(Intercept)'} | {'std'} | 4.5929 | 3.7091 | 5.6873 |

Group: Rat (6 Levels)

| Name1 | Name2 | Type | Estimate | Lower | Upper |
|---|---|---|---|---|---|
| {'(Intercept)'} | {'(Intercept)'} | {'std'} | 5.4543e-07 | NaN | NaN |

Group: Error

| Name | Estimate | Lower | Upper |
|---|---|---|---|
| {'Res Std'} | 2.3055 | 2.0969 | 2.5349 |

### Fig 2A - LME Model For Xylocaine blocker experiment

**Linear mixed-effects model fit by ML**

**Model information:**

| | |
|---|---|
| Number of observations | 117 |
| Fixed effects coefficients | 3 |
| Random effects coefficients | 41 |
| Covariance parameters | 3 |

**Formula:**

SPK ~ 1 + Condition + (1 | Neuron) + (1 | Rat)

**Model fit statistics:**

| AIC | BIC | LogLikelihood | Deviance |
|---|---|---|---|
| 543.01 | 559.59 | -265.51 | 531.01 |

**Fixed effects coefficients (95% CIs):**

| Name | Estimate | SE | tStat | DF | pValue | Lower | Upper |
|---|---|---|---|---|---|---|---|
| {'(Intercept)' } | 3.4355 | 1.3391 | 2.5655 | 114 | 0.011602 | 0.78268 | 6.0883 |
| {'Condition_pre' } | 0.17805 | 0.29649 | 0.60052 | 114 | 0.54935 | -0.40929 | 0.76538 |
| {'Condition_post'} | 0.07875 | 0.29649 | 0.26561 | 114 | 0.79102 | -0.50859 | 0.66609 |

**Random effects covariance parameters (95% CIs):**

Group: Neuron (39 Levels)

| Name1 | Name2 | Type | Estimate | Lower | Upper |
|---|---|---|---|---|---|
| {'(Intercept)'} | {'(Intercept)'} | {'std'} | 4.1057 | 3.2432 | 5.1975 |

Group: Rat (2 Levels)

| Name1 | Name2 | Type | Estimate | Lower | Upper |
|---|---|---|---|---|---|
| {'(Intercept)'} | {'(Intercept)'} | {'std'} | 1.5981 | 0.40392 | 6.323 |

Group: Error

| Name | Estimate | Lower | Upper |
|---|---|---|---|
| {'Res Std'} | 1.3092 | 1.1191 | 1.5317 |

### Fig 2B - LME Model For Clonidine blocker experiment

**Linear mixed-effects model fit by ML**

**Model information:**

| | |
|---|---|
| Number of observations | 144 |
| Fixed effects coefficients | 3 |
| Random effects coefficients | 53 |
| Covariance parameters | 3 |

**Formula:**

SPK ~ 1 + Condition + (1 | Neuron) + (1 | Rat)

**Model fit statistics:**

| AIC | BIC | LogLikelihood | Deviance |
|---|---|---|---|
| 720.51 | 738.33 | -354.25 | 708.51 |

**Fixed effects coefficients (95% CIs):**

| Name | Estimate | SE | tStat | DF | pValue | Lower | Upper |
|---|---|---|---|---|---|---|---|
| {'(Intercept)' } | 4.2279 | 0.66009 | 6.405 | 141 | 2.0945e-09 | 2.9229 | 5.5328 |
| {'Condition_pre' } | -0.12863 | 0.3546 | -0.36276 | 141 | 0.71733 | -0.82966 | 0.57239 |
| {'Condition_post'} | -0.14567 | 0.3546 | -0.4108 | 141 | 0.68184 | -0.84669 | 0.55535 |

**Random effects covariance parameters (95% CIs):**

Group: Neuron (48 Levels)

| Name1 | Name2 | Type | Estimate | Lower | Upper |
|---|---|---|---|---|---|
| {'(Intercept)'} | {'(Intercept)'} | {'std'} | 4.2305 | 3.4242 | 5.2265 |

Group: Rat (5 Levels)

| Name1 | Name2 | Type | Estimate | Lower | Upper |
|---|---|---|---|---|---|
| {'(Intercept)'} | {'(Intercept)'} | {'std'} | 1.023e-06 | NaN | NaN |

Group: Error

| Name | Estimate | Lower | Upper |
|---|---|---|---|
| {'Res Std'} | 1.7372 | 1.508 | 2.0011 |

### Fig. 3 - LME Model For All Single Units Over Time

```
Linear mixed-effects model fit by ML

Model information:
    Number of observations             616
    Fixed effects coefficients           7
    Random effects coefficients         93
    Covariance parameters                3

Formula:
    SPK ~ 1 + Condition + (1 | Neuron) + (1 | Rat)

Model fit statistics:
    AIC       BIC       LogLikelihood    Deviance
    3692.4    3736.7    -1836.2          3672.4

Fixed effects coefficients (95% CIs):
    Name                           Estimate    SE         tStat       DF     pValue        Lower       Upper
    {'(Intercept)'     }             4.2162    0.81257      5.1887    609    2.8881e-07      2.6204      5.8119
    {'Condition_dur'   }            0.56499    0.57681      0.9795    609       0.32772     -0.5678      1.6978
    {'Condition_post'  }            0.78173    0.57681      1.3553    609       0.17584    -0.35106      1.9145
    {'Condition_post15'}              1.577    0.57681       2.734    609     0.0064382     0.44425      2.7098
    {'Condition_post30'}             1.1568    0.57681      2.0054    609      0.045361    0.023964      2.2895
    {'Condition_post45'}            0.19117    0.57681     0.33143    609       0.74044    -0.94162       1.324
    {'Condition_post60'}           -0.56827    0.57681    -0.98518    609       0.32493     -1.7011     0.56452

Random effects covariance parameters (95% CIs):
Group: Neuron (88 Levels)
    Name1                  Name2                  Type           Estimate    Lower     Upper
    {'(Intercept)'}        {'(Intercept)'}        {'std'}        6.5927      5.6468    7.6971

Group: Rat (5 Levels)
    Name1                  Name2                  Type           Estimate      Lower    Upper
    {'(Intercept)'}        {'(Intercept)'}        {'std'}        6.2751e-08    NaN      NaN

Group: Error
    Name               Estimate    Lower     Upper
    {'Res Std'}        3.8262      3.6022    4.064
```

### Fig. 4 - LME Model STA Theta

```
Linear mixed-effects model fit by ML

Model information:
    Number of observations             300
    Fixed effects coefficients           3
    Random effects coefficients        106
    Covariance parameters                3

Formula:
    STA ~ 1 + Condition + (1 | Neuron) + (1 | Rat)

Model fit statistics:
    AIC        BIC        LogLikelihood    Deviance
    -1839.7    -1817.5    925.85           -1851.7

Fixed effects coefficients (95% CIs):
    Name                         Estimate      SE           tStat      DF     pValue       Lower         Upper
    {'(Intercept)'   }              1.0074    0.0016755     601.28    297            0        1.0041        1.0107
    {'Condition_pre' }          -0.0048808    0.0011429    -4.2706    297    2.627e-05      -0.00713    -0.0026316
    {'Condition_post'}          -0.0016949    0.0011429     -1.483    297      0.13913    -0.0039441    0.00055428

Random effects covariance parameters (95% CIs):
Group: Neuron (100 Levels)
    Name1                  Name2                  Type           Estimate    Lower        Upper
    {'(Intercept)'}        {'(Intercept)'}        {'std'}        0.010804    0.0091235    0.012793

Group: Rat (6 Levels)
    Name1                  Name2                  Type           Estimate     Lower         Upper
    {'(Intercept)'}        {'(Intercept)'}        {'std'}        0.0022819    0.00061823    0.0084223

Group: Error
    Name               Estimate     Lower       Upper
    {'Res Std'}        0.0080814    0.007327    0.0089135
```

### Fig. 4 - LME Model STA Gamma

```
Linear mixed-effects model fit by ML

Model information:
    Number of observations             300
    Fixed effects coefficients           3
    Random effects coefficients        106
    Covariance parameters                3

Formula:
    STA ~ 1 + Condition + (1 | Neuron) + (1 | Rat)

Model fit statistics:
    AIC      BIC        LogLikelihood    Deviance
    -3723    -3700.8    1867.5           -3735

Fixed effects coefficients (95% CIs):
    Name                         Estimate       SE            tStat       DF     pValue     Lower          Upper
    {'(Intercept)'   }              1.0007     7.951e-05       12586    297          0         1.0005        1.0008
    {'Condition_pre' }         -5.3921e-05    4.2114e-05     -1.2804    297    0.20142     -0.0001368    2.8959e-05
    {'Condition_post'}            -2.03e-05    4.2114e-05    -0.48202    297    0.63014    -0.00010318     6.258e-05

Random effects covariance parameters (95% CIs):
Group: Neuron (100 Levels)
    Name1                  Name2                  Type           Estimate      Lower         Upper
    {'(Intercept)'}        {'(Intercept)'}        {'std'}        0.00069202    0.00059515    0.00080466

Group: Rat (6 Levels)
    Name1                  Name2                  Type           Estimate     Lower       Upper
    {'(Intercept)'}        {'(Intercept)'}        {'std'}        5.731e-05    2.85e-07    0.011525

Group: Error
    Name               Estimate      Lower         Upper
    {'Res Std'}        0.00029779    0.00026999    0.00032845
```

### Fig. 4 - LME Model SFC Theta

**Linear mixed-effects model fit by ML**

**Model information:**

| | |
|---|---|
| Number of observations | 273 |
| Fixed effects coefficients | 3 |
| Random effects coefficients | 97 |
| Covariance parameters | 3 |

**Formula:**

SFC ~ 1 + Condition + (1 | Neuron) + (1 | Rat)

**Model fit statistics:**

| AIC | BIC | LogLikelihood | Deviance |
|---|---|---|---|
| -882.33 | -860.67 | 447.17 | -894.33 |

**Fixed effects coefficients (95% CIs):**

| Name | Estimate | SE | tStat | DF | pValue | Lower | Upper |
|---|---|---|---|---|---|---|---|
| {'(Intercept)' } | 1.045 | 0.011264 | 92.771 | 270 | 8.2143e-207 | 1.0228 | 1.0672 |
| {'Condition_pre' } | -0.01188 | 0.0040165 | -2.9578 | 270 | 0.0033728 | -0.019787 | -0.0039723 |
| {'Condition_post'} | -0.013398 | 0.0040165 | -3.3358 | 270 | 0.00096976 | -0.021306 | -0.0054907 |

**Random effects covariance parameters (95% CIs):**

Group: Neuron (91 Levels)

| Name1 | Name2 | Type | Estimate | Lower | Upper |
|---|---|---|---|---|---|
| {'(Intercept)'} | {'(Intercept)'} | {'std'} | 0.079029 | 0.067678 | 0.092284 |

Group: Rat (6 Levels)

| Name1 | Name2 | Type | Estimate | Lower | Upper |
|---|---|---|---|---|---|
| {'(Intercept)'} | {'(Intercept)'} | {'std'} | 0.016427 | 0.0042201 | 0.063943 |

Group: Error

| Name | Estimate | Lower | Upper |
|---|---|---|---|
| {'Res Std'} | 0.027093 | 0.024447 | 0.030024 |

### Fig. 4 - LME Model SFC Gamma

**Linear mixed-effects model fit by ML**

**Model information:**

| | |
|---|---|
| Number of observations | 273 |
| Fixed effects coefficients | 3 |
| Random effects coefficients | 97 |
| Covariance parameters | 3 |

**Formula:**

SFC ~ 1 + Condition + (1 | Neuron) + (1 | Rat)

**Model fit statistics:**

| AIC | BIC | LogLikelihood | Deviance |
|---|---|---|---|
| -1627.4 | -1605.8 | 819.72 | -1639.4 |

**Fixed effects coefficients (95% CIs):**

| Name | Estimate | SE | tStat | DF | pValue | Lower | Upper |
|---|---|---|---|---|---|---|---|
| {'(Intercept)' } | 1.0222 | 0.0040296 | 253.67 | 270 | 3.9525e-323 | 1.0142 | 1.0301 |
| {'Condition_pre' } | -0.0026411 | 0.00091372 | -2.8905 | 270 | 0.0041582 | -0.0044401 | -0.00084223 |
| {'Condition_post'} | -0.0029416 | 0.00091372 | -3.2193 | 270 | 0.0014419 | -0.0047405 | -0.0011427 |

**Random effects covariance parameters (95% CIs):**

Group: Neuron (91 Levels)

| Name1 | Name2 | Type | Estimate | Lower | Upper |
|---|---|---|---|---|---|
| {'(Intercept)'} | {'(Intercept)'} | {'std'} | 0.025505 | 0.021864 | 0.029754 |

Group: Rat (6 Levels)

| Name1 | Name2 | Type | Estimate | Lower | Upper |
|---|---|---|---|---|---|
| {'(Intercept)'} | {'(Intercept)'} | {'std'} | 0.0068874 | 0.0019948 | 0.02378 |

Group: Error

| Name | Estimate | Lower | Upper |
|---|---|---|---|
| {'Res Std'} | 0.0061634 | 0.0055617 | 0.0068302 |

### Fig. 4 - LME Model STA Clonidine Theta

**Linear mixed-effects model fit by ML**

<strong>Model information:</strong>

| | |
|---|---|
| Number of observations | 144 |
| Fixed effects coefficients | 3 |
| Random effects coefficients | 53 |
| Covariance parameters | 3 |

**Formula:**

STA ~ 1 + Condition + (1 | Neuron) + (1 | Rat)

**Model fit statistics:**

| AIC | BIC | LogLikelihood | Deviance |
|---|---|---|---|
| -1109.3 | -1091.4 | 560.63 | -1121.3 |

**Fixed effects coefficients (95% CIs):**

| Name | Estimate | SE | tStat | DF | pValue | Lower | Upper |
|---|---|---|---|---|---|---|---|
| {'(Intercept)' } | 1.0028 | 0.00080364 | 1247.8 | 141 | 6.1106e-287 | 1.0012 | 1.0044 |
| {'Condition_pre' } | -0.00052673 | 0.00079427 | -0.66316 | 141 | 0.50831 | -0.002097 | 0.0010435 |
| {'Condition_post'} | -0.00031119 | 0.00079427 | -0.39179 | 141 | 0.6958 | -0.0018814 | 0.001259 |

**Random effects covariance parameters (95% CIs):**

Group: Neuron (48 Levels)

| Name1 | Name2 | Type | Estimate | Lower | Upper |
|---|---|---|---|---|---|
| {'(Intercept)'} | {'(Intercept)'} | {'std'} | 0.0039824 | 0.0030477 | 0.0052038 |

Group: Rat (5 Levels)

| Name1 | Name2 | Type | Estimate | Lower | Upper |
|---|---|---|---|---|---|
| {'(Intercept)'} | {'(Intercept)'} | {'std'} | 7.1687e-13 | NaN | NaN |

Group: Error

| Name | Estimate | Lower | Upper |
|---|---|---|---|
| {'Res Std'} | 0.0038911 | 0.0033779 | 0.0044824 |

### Fig. 4 - LME Model STA Clonidine Gamma

**Linear mixed-effects model fit by ML**

**Model information:**

| | |
|---|---|
| Number of observations | 144 |
| Fixed effects coefficients | 3 |
| Random effects coefficients | 53 |
| Covariance parameters | 3 |

**Formula:**

STA ~ 1 + Condition + (1 | Neuron) + (1 | Rat)

**Model fit statistics:**

| AIC | BIC | LogLikelihood | Deviance |
|---|---|---|---|
| -2084.3 | -2066.5 | 1048.2 | -2096.3 |

**Fixed effects coefficients (95% CIs):**

| Name | Estimate | SE | tStat | DF | pValue | Lower | Upper |
|---|---|---|---|---|---|---|---|
| {'(Intercept)' } | 1.0003 | 5.1255e-05 | 19517 | 141 | 0 | 1.0002 | 1.0004 |
| {'Condition_pre' } | -2.9931e-05 | 1.7942e-05 | -1.6683 | 141 | 0.097484 | -6.5401e-05 | 5.5381e-06 |
| {'Condition_post'} | -2.9945e-06 | 1.7942e-05 | -0.1669 | 141 | 0.86769 | -3.8464e-05 | 3.2475e-05 |

**Random effects covariance parameters (95% CIs):**

Group: Neuron (48 Levels)

| Name1 | Name2 | Type | Estimate | Lower | Upper |
|---|---|---|---|---|---|
| {'(Intercept)'} | {'(Intercept)'} | {'std'} | 0.00034406 | 0.00028045 | 0.00042209 |

Group: Rat (5 Levels)

| Name1 | Name2 | Type | Estimate | Lower | Upper |
|---|---|---|---|---|---|
| {'(Intercept)'} | {'(Intercept)'} | {'std'} | 4.457e-12 | NaN | NaN |

Group: Error

| Name | Estimate | Lower | Upper |
|---|---|---|---|
| {'Res Std'} | 8.7896e-05 | 7.6302e-05 | 0.00010125 |

### Fig. 4 - LME Model SFC Clonidine Theta

```
Linear mixed-effects model fit by ML

Model information:
    Number of observations            117
    Fixed effects coefficients          3
    Random effects coefficients        44
    Covariance parameters               3

Formula:
    SFC ~ 1 + Condition + (1 | Neuron) + (1 | Rat)

Model fit statistics:
    AIC        BIC        LogLikelihood    Deviance
    -460.15    -443.57    236.07           -472.15

Fixed effects coefficients (95% CIs):
    Name                        Estimate     SE           tStat     DF     pValue        Lower          Upper
    {'(Intercept)'   }             1.0203     0.011493    88.773    114    4.564e-107       0.99752       1.0431
    {'Condition_pre' }          0.0072907    0.0055382    1.3165    114       0.19066    -0.0036803     0.018262
    {'Condition_post'}            0.01004    0.0055382    1.8128    114      0.072489   -0.00093131     0.021011

Random effects covariance parameters (95% CIs):
Group: Neuron (39 Levels)
    Name1                  Name2                  Type           Estimate    Lower       Upper
    {'(Intercept)'}        {'(Intercept)'}        {'std'}        0.025393    0.018331    0.035177

Group: Rat (5 Levels)
    Name1                  Name2                  Type           Estimate    Lower        Upper
    {'(Intercept)'}        {'(Intercept)'}        {'std'}        0.021688    0.0080185    0.05866

Group: Error
    Name               Estimate    Lower       Upper
    {'Res Std'}        0.024456    0.020904    0.028611
```

### Fig. 4 - LME Model SFC Clonidine Gamma

```
Linear mixed-effects model fit by ML

Model information:
    Number of observations            117
    Fixed effects coefficients          3
    Random effects coefficients        44
    Covariance parameters               3

<strong>Formula:</strong>
    SFC ~ 1 + Condition + (1 | Neuron) + (1 | Rat)

Model fit statistics:
    AIC        BIC       LogLikelihood    Deviance
    -371.87    -355.3    191.94           -383.87

Fixed effects coefficients (95% CIs):
    Name                         Estimate      SE           tStat       DF     pValue         Lower        Upper
    {'(Intercept)'   }              1.0127    0.0075125       134.8    114    1.5045e-127      0.99781       1.0276
    {'Condition_pre' }          -0.0018511     0.010624    -0.17423    114        0.86199    -0.022898     0.019195
    {'Condition_post'}           0.0078536     0.010624     0.73922    114        0.46129    -0.013193       0.0289

Random effects covariance parameters (95% CIs):
Group: Neuron (39 Levels)
    Name1                  Name2                  Type           Estimate      Lower    Upper
    {'(Intercept)'}        {'(Intercept)'}        {'std'}        9.8392e-09    NaN      NaN

Group: Rat (5 Levels)
    Name1                  Name2                  Type           Estimate      Lower    Upper
    {'(Intercept)'}        {'(Intercept)'}        {'std'}        7.1823e-10    NaN      NaN

Group: Error
    Name               Estimate    Lower       Upper
    {'Res Std'}        0.046915    0.041273    0.053328
```
